# miR-155 expression and function in leukemic stem cells

**DOI:** 10.1101/2024.02.29.582835

**Authors:** Ramiro Garzon, Carlo M Croce

## Abstract

It has been reported that elevated levels of miR-155 have been reported in patients with Acute Myeloid Leukemia (AML) bearing *FLT3*-ITD mutations and is independent of *FLT3*-ITD signaling. However, it is unclear how miR-155 is expressed in leukemic stem cells (LCS) and whether miR-155 has any role regulating LSC functions. In this manuscript, we defined the expression of miR-155 in clearly defined LCSs population and showed that miR-155 is regulating self-renewal and quiescence of LSCs.

## INTRODUCTION

Acute myeloid leukemia (AML) is a devastating malignant disease of the hematopoietic system that leads initially to bone marrow (BM) failure and to death if left untreated. Despite recent advances in our understanding of AML biology and the use of intensive treatments, the prognosis remains poor with only 30-40% of younger (<60 years) and <10% of older (≥60 years) adult AML patients achieving long-term overall survival^1,2^. Thus, novel therapies targeting key leukemogenic pathways in AML are needed.

MicroRNAs (miRs) are small non-coding RNAs that control the expression of proteins involved in cellular homeostatic mechanisms^3^. MiRs are dysregulated and/or mutated in AML and are known to play a key role in leukemogenesis^4^. Among microRNAs, miR-155 has emerged as one of the most frequently dysregulated oncomiRs in cancer and leukemia and has been invariably associated with worse outcome^4-8^. In murine models, sustained overexpression of miR-155 in B cells results in an aggressive pre-B cell leukemia^6^.

In our AML studies we found that: 1) higher expression of miR-155 is associated with worse outcome^7^; 2) higher expression of miR-155 is associated with *FLT3* internal tandem duplication (*FLT3*-ITD), a biomarker for poor prognosis^8^ and 3) miR-155 upregulation is mechanistically and prognostically independent from *FLT3*-ITD which may represent a second “hit” in *FLT3*-ITD-positive clonal cells^7-8^.

Leukemic stem cells (LSC) represent a relatively small population of leukemic cells that have acquired abnormal self-renewal and partial maturation ability^9^. These cells give rise to the bulk population of more mature blasts lacking self-renewal capacity. The general view is that LSCs are responsible for disease initiation and maintenance in AML^9^. Thus, targeting LSCs represents an essential step for complete eradication of the disease. However, LSCs are resistant to conventional chemotherapy regimens, and novel approaches are needed to eliminate these cells to improve clinical outcome. Identification of LSC-enriched cell subpopulation requires the use of complementary immunophenotypic and functional assays. Our group reported high expression of miR-155 to be associated with an LSC-gene expression signature predictive of poor outcome in AML^10^. This data led us to hypothesize a role for miR-155 in LSC biology. Here, we are aiming to understand the impact of miR-155 in leukemic stem cells functions.

## METHODS

**Northern blotting** of *miR*-*155* in MV4-11 cells incubated for 24 and 48 h with SU14813 (300 nM) or DMSO (vehicle control). Total RNA was obtained with TRIzol, and Northern blotting was performed with a probe antisense to *miR*-*155*. The blot was stripped and reprobed with U6 for loading control. Northern blotting conditions as described in detail elsewhere^8^.

### Long-term colony initiating cells (LT-ICS) assays

Briefly, CD34+CD38- and CD34+CD38+ subpopulations were isolated using flow sorting and then serial dilutions of each of the two subpopulations were seeded onto irradiated stromal layers. After 6-weeks of culture, cobblestone area forming cells (CAFCs) were scored to determine the presence of LTC-ICs. The frequency of LSCs was calculated using Poisson statistics with the L-Calc Software for limiting dilution analysis (StemCell Technologies)

**CFSE (CellTrace CFSE Proliferation Kit**) according to the manufacturer’s instructions and treated with -antimiR-155 or SCR dosed three days after staining.

## RESULTS

### Mir-155 expression in leukemia cells lines after FLT3-ITD inhibition

It has been reported that high levels of miR-155 have been reported in patients with AML bearing *FLT3*-ITD mutations^7-8^. Our group previously reported that miR-155 levels measured by Northern Blotting in MV-4-11 cell lines did not change after treatment in vitro with the *FLT3*-ITD inhibitor SU14813^8^. Here we repeated that experiment to confirm these results. As seen in Figure 1, miR-155 levels were unchanged after therapy with SU14813. Raw data is in supplemental Fig. 1 and 2.

**Figure 1:**
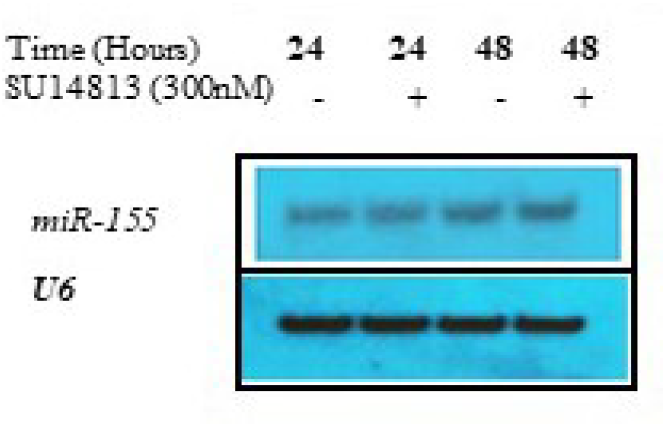
Northern blotting of *miR*-*155* in MV4-11 cells incubated for 24 and 48 h with SU14813 (300 nM)or DMSO (vehicle control).

### Mir-155 expression in primary AML samples LCSs

To understand the expression of miR-155 in in LSC-enriched cell populations in primary AML patient samples, we must first identify which CD34/CD38 compartment (CD34+CD38-vs. CD34+CD38+) is enriched for LSCs in each patient sample by measuring the frequency of LTC-ICs using limiting dilution assays (LDA). Briefly, CD34+CD38- and CD34+CD38+ subpopulations were isolated using flow sorting and then serial dilutions of each of the two subpopulations were seeded onto irradiated stromal layers. After 6-weeks of culture, cobblestone area forming cells (CAFCs) were scored to determine the presence of LTC-ICs. We performed LTC-ICs from 4 patients with cytogenetically normal (CN) AML (two with *FLT3*-ITD+ and two with *FLT3*-wild type) and from 3 cord blood (CB) from healthy donors. MiR-155 levels were measured in AML and CB samples by real time RT-PCR and results were normalized to an endogenous reference (U44) and reported relative to the levels in pooled normal CB HSC (CB+-). As shown in Figure 2, miR-155 expression was increased in *FLT3*-ITD+ CD34+CD38- and CD34+CD38+ populations with respect to normal CBs and CN-AML cases with *FLT3*-wildtype. In addition, miR-155 was higher expressed in more primitive cell populations according to the LT-ICS assays that showed enrichment with LSC.

**Figure 2.**
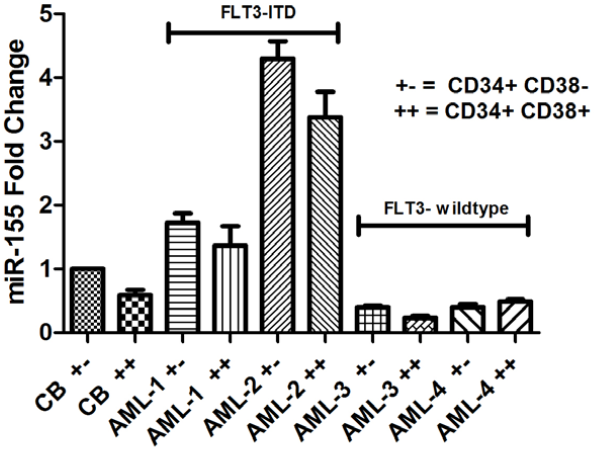
miR-155 expression in CN-AML progenitors.

### Functional implications of miR-155 in LSC

To gain insights about the functional role of miR-155 in regulating LSC functions, we then performed knock-down of miR-155 using LNA-antimiR-155 or scrambled controls (SCR) in CD34+ sorted cells from two primary *FLT3*-ITD+ AML samples and a CB (Control) for 24 hours and performed colony forming units (CFUs) assays. Primary CFU cells were harvested on day fourteen then used in re-plating assays to assess their self-renewal capacity. As shown in Figure 3, CD34+ cells treated with antimiR-155 showed a significant decrease in the number of colonies with respect to SCR-treated controls after replating. Of interest is that there was no statistical difference between LNA antimiR-155 or control treated CB cells, suggesting that the miR-treatment would not impact on the self-renewal function of normal HSC. Altogether, our data support the hypothesis that miR-155 plays a role in LSC self-renewal.

**Figure 3.**
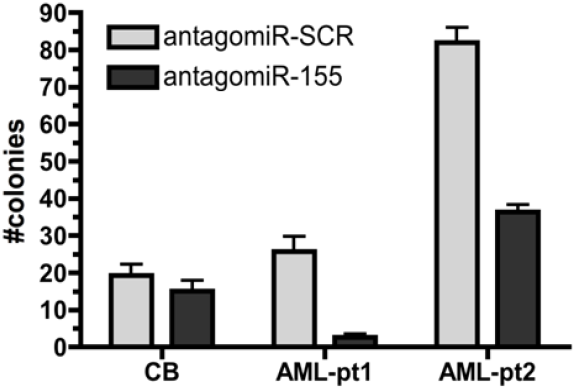
Role of miR-155 in AML self-renewal.

Next, to determine the impact of miR-155 in the LSC frequency we performed LTC-IC assays in primary FLT3-ITD+ CD34^+^ patients’ samples (N=3) after miR-155 knock down using nanoparticle transferrin conjugated antimiR-155 or scrambled mixmers. Indeed, we found that miR-155 knock-down decreased the LSC frequency compared to SCR (P=.001, P=.007, P=.002, respectively).

To determine changes within the quiescent cell subpopulation, CD34+ AML cells from three patient samples were stained with CFSE (CellTrace CFSE Proliferation Kit) according to the manufacturer’s instructions and treated with antimiR-155 or SCR dosed three days after staining. After 3 further days in culture cells were harvested and stained with anti-CD34 PEcy7, viability stain DAPI. Cytokine-deprived non-dividing CD34+ AML cells were used as a control to identify the CFSE fluorescent quiescent population. Quiescent cells (CFSEmaxCD34+) were reported as a fraction of the initial number of CD34+ cells. In all analyzed patient samples, we found a significant decrease in the number of CTV^high^/CD34^+^ cells after treatment with antimiR-155 compared to control (*P*<.01, *P*=.022, *P*=.012, respectively), indicating that miR-155 regulates LSC quiescence in AML.

## DISCUSSION

MiR-155 is hosted at the *BIC* gene locus on chromosome 21 and is one of the most deregulated miRs in cancer^4-8^. High miR-155 expression is associated with aggressive phenotypes and worse outcomes in solid tumors, lymphomas, and leukemia^4-8^. Although miR-155 is dispensable for normal lymphocyte and myeloid development, it plays a crucial role in the immune system^11-12^. miR-155 is also induced in hematopoietic stem cells (HSCs), progenitors and myeloid cells during inflammatory responses under the transcriptional control of NF-κB and AP-1^13-15^. In myeloid cells, among other targets, miR-155 targets two important genes involved in hematopoiesis and inflammation: 1) Src homology-2 domain-containing inositol 5-phosphatase 1 (SHIP1) a phosphatase involved in PI3K signaling pathway implicated in cell proliferation and survival, and 2) PU.1 a transcription factor implicated in myeloid differentiation^16-17^. This work led to the hypothesis that deregulated miR-155 expression contributes to abnormal proliferation, survival, and maturation arrest of myeloid blasts and potentially to the development of a leukemia phenotype. Although we showed that sustained upregulation of miR-155 in mouse lymphocyte precursors is sufficient to induce aggressive acute leukemia^6^, transient miR-155 overexpression in normal murine HSC resulted in aberrant myeloproliferation without leukemic transformation, suggesting that deregulated miR-155 requires additional ‘hits’ to progress to overt AML^17^.

Among the numerous recurrent gene mutations occurring in AML, *FLT3* mutations are one of the most common^1-2^. *FLT3* is a member of the class III receptor tyrosine kinases involved in HSC/progenitors biology. This gene is mutated in 25-30% of AML patients, either by internal tandem duplications (ITD) of the juxtamembrane domain or by point mutations usually involving the kinase domain^18^. Both types of mutations constitutively activate *FLT3*, but only *FLT3-*ITD has been invariably associated with poor prognosis^18-20^. In addition, constitutively activated *FLT3* is a therapeutic target for tyrosine kinase inhibitors (TKIs). These data along with the fact that *FLT3-*ITD overexpression in mice was shown to induce uncontrolled myeloproliferation and not AML^21^, support the view that additional ‘hits’ must be acquired for the development of overt leukemia phenotype in *FLT3* mutated blasts.

Here, we reported that miR-155 is highly expressed in *FLT3*-ITD positive blasts, miR-155 is highly expressed in both CD34+CD38- and CD34+CD38+ subpopulations and that blocking miR-155 negatively impacts on self-renewal and quiescence suggesting an active role of miR-155 in LSC activity.

Elucidating the functional role of miR-155 in LSC biology will create the path by which novel approaches designed to eradicate LSCs and target the initiating clones (the cause of most leukemia relapses) can be developed.

## Supplemental Figures

**Figure S1.**
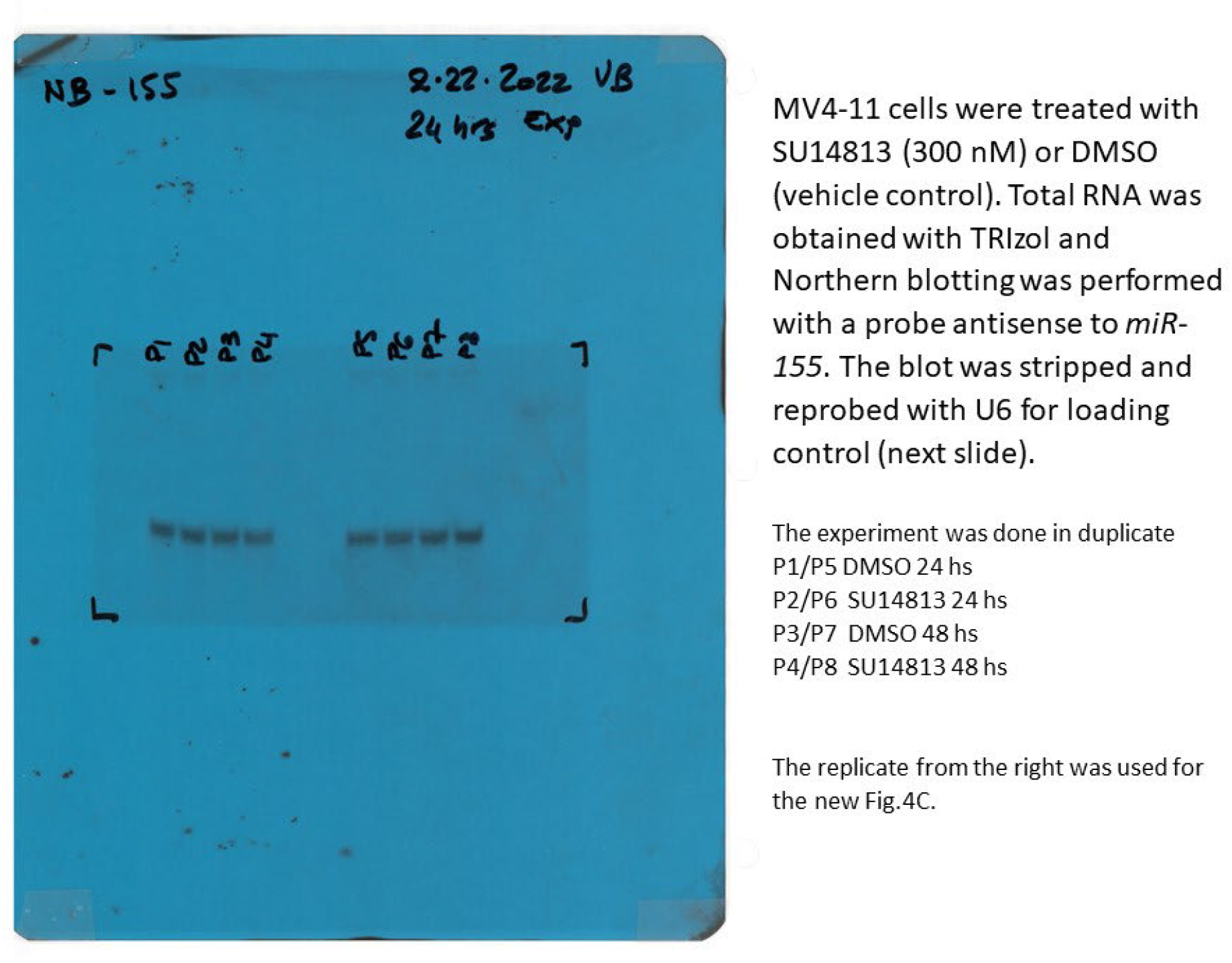

**Figure S2.**
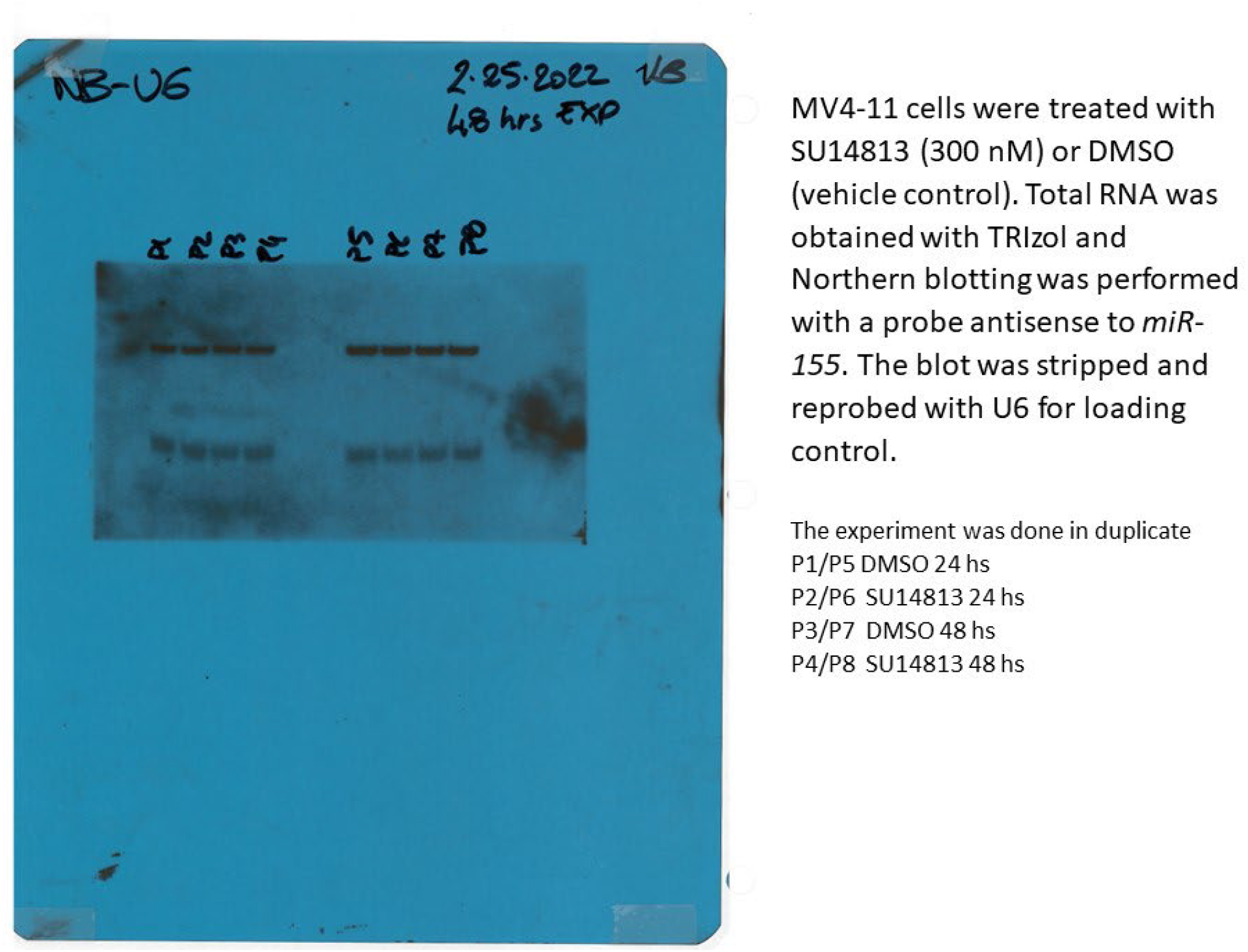

