## Supplementary figures and images for "miR-155 expression and function in leukemic stem cells"

### Figure S1

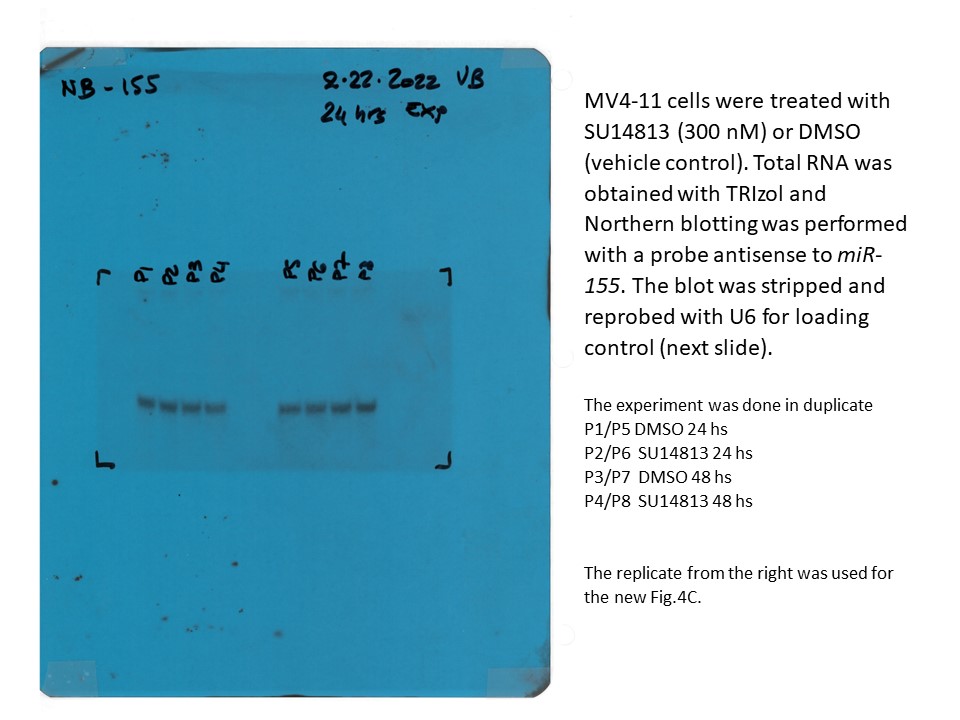

### Figure S 2

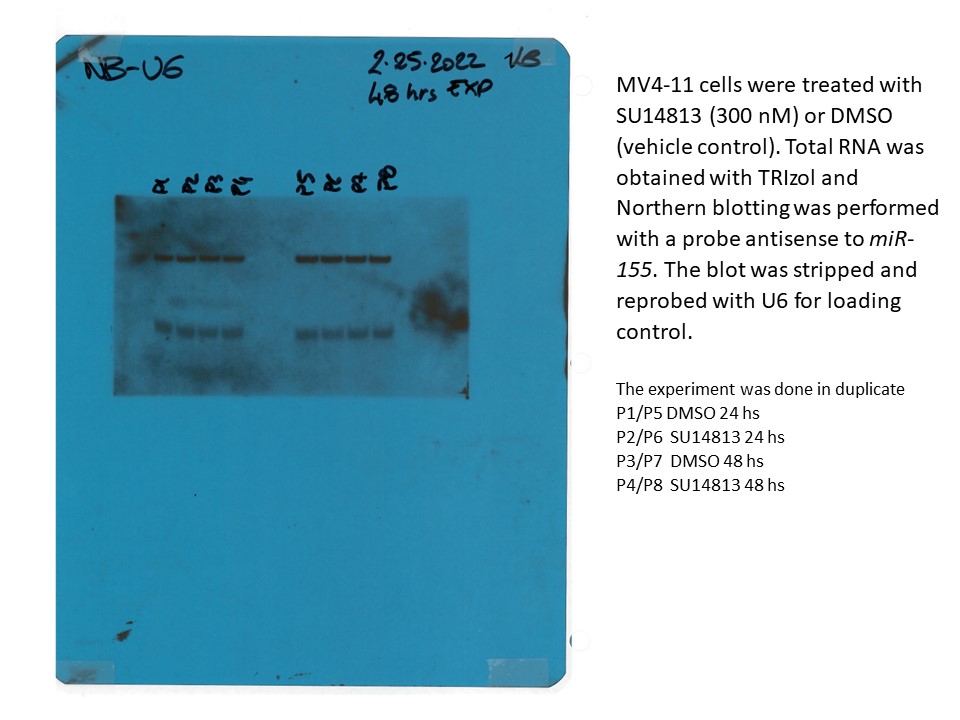
